# Sea slugs unplugged: Review of subpopulation sampling in phylogeographic studies of nudibranchs and their allies

**DOI:** 10.64898/2026.10.29.762307

**Authors:** Michael W. Hart, Miranda T. Dennis

## Abstract

Nudibranchs and other sea slugs include many cryptic species groups with unknown phenotypic and ecological diversity. How broadly and deeply should researchers sample those cryptic species pairs to capture the genetic variation within each species and potential adaptive differences between species? Fast and inexpensive surveys of genetic diversity that sample a single well-known locus across the geographical range (“phylogeography unplugged”) have long been used to identify structured populations and guide choice of sampling localities for subsequent comparative genomic or phenotypic studies of the same species. Results from such studies might also be used to better understand the expected relationships between sampling and population structure in future comparative studies in other species (including many marine gastropods) for which an unplugged phylogeographic survey is not available or impractical. We reviewed analyses of molecular variance that used mitochondrial barcode sequences from marine sea slugs, and we asked whether there are characteristic patterns of subpopulation sampling that are associated with the magnitude and geographical scale of differentiation. We found wide variation in sampling designs: studies with both many and few samples discovered significant population differentiation or broadly similar distribution of haplotypes on both large and small spatial scales, in species from different higher taxonomic groups and oceans. We found two significant associations between sampling variables and population genetic variables, and we suggest using those associations to help guide (or understand the consequences of) sampling design in future studies.

---

Genome-wide analyses of genetic variation in molluscs and other animals are important sources of insight into phylogenetic relationships among species, demographic history within species, causes of or limits on the evolution of adaptations and adaptive differences, and conservation risk or triage (Wood et al. 2008; Kocot et al. 2011; Servedio et al. 2011; Feder et al. 2012; Ravinet et al. 2016; Hohenlohe et al. 2021). The return on investment in a genome-wide analysis can be improved by first sampling spatial variation among subpopulations across the species range to predict patterns of structure that might be expected in genome-wide analyses (Saavedra and Peña 2005; Vendrami et al. 2019). Since the 21^st^-century revolution in genome-wide sequencing and bioinformatics there remains an important place for preliminary surveys of range-wide geographic variation using a single locus with well-known genetic and molecular properties as a starting point (Avise et al. 1987; Avise 2000; Garrick et al. 2015). Bowen et al. (2014) called this approach “phylogeography unplugged”.

Many marine gastropods are small, ephemeral, scarce, or otherwise challenging to collect (Jörger et al. 2012, 2014) and few of these have been the target of an unplugged phylogeographic analysis (e.g., Maroni et al. 2022). Despite that gap in knowledge, such species can provide important and useful opportunities for genome-wide study of variation and adaptation. Examples include cryptic species groups of nudibranchs and other sea slugs (e.g., Lindsay and Valdés 2016; Lindsay et al. 2016; Jung et al. 2020), including some that have long been used as model organisms in neuroscience (Katz & Quinlan 2019) and natural products chemistry (Dean and Prinsep 2017). Phenotypic and genomic variation represented by cryptic species may be an important missing component of documented biodiversity in such taxa (Struck et al. 2017; Shin and Allmon 2023). Unfortunately, our ignorance of geographic distributions of cryptic species (Bickford et al. 2006) and the extent and depth of genetic structure among subpopulations within cryptic species, combined with practical obstacles to carrying out an unplugged survey across their possible geographic ranges, are significant barriers to better understanding of adaptive divergence among cryptic species.

In the absence of an unplugged survey of far-flung or scarce subpopulations in inaccessible habitats, how intensively should researchers sample subpopulations and individuals to capture possible phylogeographical structure and coalescent depth in any new study of genome-wide variation? Adequate sampling is important for the design of genome-wide studies (Nazareno et al. 2017; Aguirre-Liguori et al. 2025), but practical limits (time, funds) may impose tradeoffs and require compromises (e.g., Felsenstein 2006). In our recent and ongoing work to characterize the demographic history of divergence among cryptic species in several sea slugs in the northeastern Pacific (Dennis et al. 2025), we lacked both detailed knowledge of their biogeography and the resources needed to first carry out systematic sampling in an unplugged survey, and we have depended instead on convenience sampling of relatively accessible subpopulations (and assumed that we captured a representative portion of their genetic diversity).

Are there other sources of information that could improve either the design of new genome-wide studies that lack an unplugged survey, or our understanding of the consequences of convenience sampling (rather than systematic range-wide sampling)? We reviewed phylogeographic studies in nudibranchs and other sea slugs that used the same genetic marker and analytical method (analysis of molecular variance). We focused mainly on variation among studies in the sampling scheme, especially the number and geographical distribution of subpopulations and individuals. We also considered one biological trait of the organisms that was emphasized in all of the reviewed studies (the mode of larval development and potential for gene flow by planktonic larval dispersal).

We used the review to address three related problems. (i) We summarize variation in sampling designs used by previous researchers. (ii) We speculate about possible reasons for choices of sampling designs. (iii) We ask whether the patterns we found in previous studies might be useful for choosing sampling designs in future studies. We found (i) wide variation in the size and geographical distribution of samples, including several correlations among sampling variables that probably reflect (ii) researcher expectations about the scale and extent of genetic variation among subpopulations, and tradeoffs or choices about how to sample organisms and subpopulations given those expectations. We also found two associations between a population genetic variable and several of the sampling variables, and (iii) cautiously interpret those associations in the context of future study design choices.

## MATERIALS AND METHODS

### Literature review and inclusion criteria

We searched the Web of Science and Google Scholar databases on 7 October 2025. In the Web of Science we searched “nudibranch*” and “genet*” in all fields (145 records). The same search terms in Google Scholar returned almost 4000 records so we used a more selective search string: "nudibranch* and ("population genetic" or "population genetics") and (differentiat* or structure* or isolat* or "gene flow")" (214 records). We combined those two lists, removed duplicate titles, and added several titles already known to us or suggested by colleagues but not included in the search results (347 records after filtering; Appendix 1).

We then reviewed each of those articles to identify studies that estimated subpopulation structure as *Φ* _ST_ values using mitochondrial *COI* sequences in analysis of molecular variance (AMOVA; Excoffier et al. 1992). The majority of articles (195) were not studies of nudibranchs or other sea slugs (Appendix 1); of the rest, 42 articles did not include population genetic data; 93 articles included population genetic data but no analysis of spatial structure using analysis of molecular variance (these articles included a mix of studies that used other data types, focused on other questions such as cryptic species identification, or lacked details about sampling and analysis); and 2 articles were abstracts only. We analyzed studies from the remaining 15 articles that included AMOVA results for 20 species (Table 1).

**Table 1.** Sampling and population genetic variables for studies of 20 sea slug species from 15 articles. Taxon names are from the World Register of Marine Species (July 2026). The numbers of subpopulations (n) and individuals (N) are those in the analysis of molecular variance. Median great circle distance and median *Φ* _ST_ value between pairs of subpopulations includes the interquartile range (IQR) for studies with 4 or more subpopulations. For two studies the smallest distance between any pair of subpopulations with a *Φ* _ST_ value significantly different from zero is marked by an asterisk and shown as 1 + the largest distance between subpopulations (because no pairwise *Φ* _ST_ values were significantly different from zero).

| Taxon | Mode of development | n | N | Median distance (IQR), km | Largest distance, km | Median *Φ*<sub>ST</sub> (IQR) | Smallest distance with *Φ*<sub>ST</sub> > 0, km | Reference |
|---|---|---|---|---|---|---|---|---|
| Aeolidina (nudibranchs) |  |  |  |  |  |  |  |  |
| *Dondice parguerensis* Brandon and Cutress, 1985 | Feeding planktonic veliger | 2 | 43 | 1899 | 1899 | 0.625 | 1899 | García-Méndez et al. (2022) |
| *Eubranchulus rupium* (Møller, 1842) | Feeding planktonic veliger | 3 | 108 | 5846 | 6048 | 0.0155 | 305 | Grishina et al. (2024), as *Eubranchus rupium* |
| *Glaucus atlanticus* Forster, 1777 | Feeding planktonic veliger | 5 | 97 | 12944 (10318–21501) | 32647 | 0.244 (0.015–0.318) | 8672 | Churchill et al. (2014); Ross and Quetin (1990) |
| *Phidiana hiltoni* (O'Donoghue, 1927) | Nonfeeding planktonic veliger | 3 | 42 | 304 | 377 | 0.660 | 304 | King et al. (2019) |
| Pleurobranchida |  |  |  |  |  |  |  |  |
| *Pleurobranchaea maculata* (Quoy and Gaimard, 1832) | Feeding planktonic veliger | 5 | 146 | 441 (154–497) | 552 | 0.0175 (0.008–0.020) | 553* | Yildirim et al. (2018) |
| Doridida |  |  |  |  |  |  |  |  |
| *Austrodoris kerguelenensis* (Bergh, 1884) species 24 | Direct development; no planktonic larva | 8 | 356 | 1012 (574–1634) | 2163 | 0.452 (0.267–0.542) | 101 | Maroni and Wilson (2022), as *Doris kerguelenensis* |
| *Austrodoris kerguelenensis* (Bergh, 1884) species 28 | Direct development; no planktonic larva | 2 | 79 | 7605 | 7605 | 0.588 | 7605 | Maroni and Wilson (2022), as *Doris kerguelenensis* |
| *Austrodoris kerguelenensis* (Bergh, 1884) species 29 | Direct development; no planktonic larva | 5 | 279 | 515 (234–689) | 1067 | 0.154 (0.075–0.210) | 119 | Maroni and Wilson (2022), as *Doris kerguelenensis* |
| *Goniobranchus splendidus* (Angas, 1864) | Nonfeeding planktonic veliger | 5 | 105 | 432 (283–681) | 847 | 0.715 (0.686–0.844) | 93 | Winters et al. (2017) |
| *Onchidoris muricata* (O. F. Müller, 1776) | Feeding planktonic veliger | 4 | 151 | 3775 (1515–6057) | 7548 | 0.56 (0.036–0.071) | 264 | Ekimova et al. (2023) |
| Aplysiida (sea hares) |  |  |  |  |  |  |  |  |
| *Aplysia dactylomela* Rang, 1828 | Feeding planktonic veliger | 2 | 19 | 8134 | 8134 | 0.160 | 8135* | Valdés et al. (2013) |
| *Phyllaplysia taylori* Dall, 1900 | Direct development; no planktonic larva | 11 | 160 | 849 (303–1540) | 1963 | 0.850 (0.340–1.0) | 133 | Louie (2003) |
| Plakobranchoidea (sacoglossans) |  |  |  |  |  |  |  |  |
| *Alderia harvardiensis* A. A. Gould, 1870 | Feeding planktonic veliger | 10 | 94 | 4097 (733–5030) | 10362 | 0.039 (-0.012–0.143) | 524 | Ellingson et al. (2025) |
| *Costasiella ocellifera* (Simroth, 1895) | Feeding planktonic veliger and direct development (polymorphic) | 12 | 151 | 740 (363–1464) | 2878 | 0.087 (0.020–0.603) | 66 | Ellingson and Krug (2015) |
| *Cyerce nicholasi* K. Moreno, Rico, Middlebrooks, Medrano, Á. Valdés and P. J. Krug, 2023 | Nonfeeding planktonic veliger | 4 | 16 | 204 (137–292) | 359 | 0.443 (0.273–0.593) | 82 | Moreno (2024) |
| *Elysia papillosa* A. E. Verrill, 1901 | Feeding and nonfeeding planktonic veliger (polymorphic) | 9 | 110 | 1448 (372–1836) | 3132 | 0.653 (0.428–0.780) | 82 | Krug et al. (2026) |
| *Elysia patina* Ev. Marcus, 1980 | Nonfeeding planktonic veliger | 3 | 21 | 250 | 418 | 0.83 | 168 | Krug et al. (2026) |
| *Elysia taino* Krug, Vendetti and Á. Valdés, 2016 | Feeding planktonic veliger | 7 | 56 | 610 (283–1575) | 2051 | 0.0815 (0.015–0.365) | 1492 | Krug et al. (2026) |
| *Elysia zuleicae* Ortea and Espinosa, 2002 | Feeding and nonfeeding planktonic veliger (polymorphic) | 13 | 155 | 1147 (443–1634) | 3132 | 0.0883 (0.021–0.130) | 137 | Krug et al. (2026) |
| Eupulmonata |  |  |  |  |  |  |  |  |
| *Onchidium reevesii* (J. E. Gray, 1850) | Feeding planktonic veliger | 8 | 202 | 670 (375–1079) | 1858 | 0.004 (0.003–0.075) | 295 | Zhou et al. (2016); Shen et al. (2006), as *Onchidium* sp. |

### Analysis

We used six quantitative variables (Table 1) to characterize each of those studies. Because many authors noted the mode of reproduction and development of their study organisms, and the potential for gene flow via planktonic dispersal of veliger larvae among subpopulations, we also noted one qualitative variable: the mode of larval development (planktonic veligers or benthic direct development).

We characterized four sampling variables in each study including (1) the number of subpopulations analyzed in the AMOVA and (2) the total number of individual sea slugs summed across all subpopulations. We characterized the spatial distribution of the study as (3) the median pairwise distance among all pairs of subpopulations and (4) the largest distance between subpopulations. The median distance among subpopulations can be thought of as the intensity or granularity of geographical sampling; the largest distance can be thought of as the geographical range of the study (which may be much smaller than the range of the species).

We characterized population genetic structure as (5) the median pairwise differentiation (*Φ* _ST_) among all subpopulations and (6) the smallest distance (km) between any pair of subpopulations that had a pairwise *Φ* _ST_ value significantly greater than zero in the AMOVA. For two studies (Valdés et al. 2014; Yildirim et al. 2018) that found no pairwise *Φ* _ST_ values significantly different from zero, we set this variable to 1 + the largest distance (km) between subpopulations. The median pairwise *Φ* _ST_ value reflects both the relative magnitude and the variation in differentiation among subpopulations; the smallest distance with a significant *Φ* _ST_ value reflects the granularity of that differentiation (large values reflect coarse spatial variation, small values reflect fine-scale variation).

Although values for variables (1)–(4) reflect in part the distribution and abundance of the organisms, we call them sampling variables because their values in each of the studies also reflect choices or tradeoffs that confronted the researchers in their sampling design for the study. Researcher decisions about how to resolve those choices or tradeoffs probably depend in part on prior knowledge or expectations about the magnitude or geographical scale of population differentiation. Sampling choices or tradeoffs might have a causal effect on the population structure that can be detected from those samples, but expectations about the magnitude or geographical scale of population structure might also have a causal effect on sampling choices. Because we could not know whether one or both of those opposite causal effects were important in any individual study (or different among studies), we describe the associations between sampling variables (1)–(4) on the one hand, and population genetic variables (5)–(6) on the other, but we try to avoid ascribing a causal effect to either of the two associations that were apparent in our results.

We obtained the sample coordinates for each specimen from each article or its supporting information files (and converted coordinates to degree decimal values where necessary). For a subpopulation that included individual organisms pooled from >1 collecting locality we calculated latitude and longitude for the subpopulation as the mean value for all the pooled localities. We used Claude Sonnet 4.2 to calculate great circle distances among all pairs of subpopulations. We tested several of the Claude calculations by repeating them using a conventional calculator (https://www.nhc.noaa.gov/gccalc.shtml) and found the results from the two methods were always within ∼1% of each other. We used those great circle distances to characterize the spatial distributions of subpopulations (rather than other distance measures that include biogeographical factors such as coastline shape or the direction and velocity of ocean currents). All pairwise distances calculated in our analysis (and associated *Φ* _ST_ values from each published article) are included in Appendix 2.

For most studies we used pairwise *Φ* _ST_ values between pairs of subpopulations taken from each article or its supporting information files. We included only subpopulation pairs analyzed by the original researchers, and excluded samples from other localities that the authors argued should be excluded (e.g., those with small sample sizes). For some studies we also excluded some subpopulations in our analyses for similar reasons (e.g., due to small sample sizes or missing geographical coordinates). All of the analyzed subpopulations and notes about inclusion or exclusion are identified in Appendix 2.

For one study (Garcia-Mendez et al., 2022) that was focused on population differentiation in the aeolid nudibranch *Dondice parguerensis* and was otherwise relevant to our review but did not include an AMOVA, we used Claude Sonnet 4.6 to calculate AMOVA coefficients including *Φ* _ST_ between subpopulations (see the prompts, responses, and script in Appendix 2). We tested the ability of Claude to correctly carry out AMOVA by analyzing two other COI sequence alignments from other studies for which we had the original data and published AMOVA results (Ekimova et al., 2023; Hart et al., 2012). In both tests Claude calculated *Φ* _ST_ values and other AMOVA coefficients within <1% of the published values.

To explore covariation among the four sampling variables and the two population genetic variables we used Claude Sonnet 4.6 to carry out a principal components analysis of the six variables for the 20 reviewed species (Table 1; see the prompts, responses, and script in Appendix 3). We used the correlations among variables and the variation among studies along the first and second principal component axes as context for understanding variation among population genetic results.

## RESULTS

We reviewed studies of 20 species including four aeolid nudibranchs, a pleurobranch, and five dorids; two sea hares; seven sacoglossans; and a pulmonate (Table 1). Nine species (including the pulmonate) develop as long-lived planktotrophic veliger larvae with potential for long-distance dispersal and gene flow in ocean currents; in six species some or all subpopulations reproduce as nonfeeding veligers with brief planktonic periods and limited dispersal potential; four other species have direct development of juveniles within a benthic egg mass and lack planktonic larval dispersal; one study (Ellingson and Krug 2015) focused on a species (*Costasiella ocellifera*) with polymorphic reproduction including both planktonic veligers in some subpopulations and direct development in others. The study organisms and subpopulations were widely distributed among habitats and oceans, including one species (*Glaucus atlanticus*) with pelagic adults and larvae (Churchill et al. 2014), and multiple species from the Southern Ocean (Maroni and Wilson 2022) and the Pacific and Atlantic Oceans.

Sampling varied from 2 to 13 subpopulations and 16 to 356 individuals per study (Table 1). The geographical distribution of subpopulations was also highly variable among studies: the largest distance between subpopulations varied from ∼300 to ∼30,000 km; the median distance among all pairs of subpopulations within a study also varied from ∼200 to ∼13,000 km (Figure 1).

**Figure 1.**
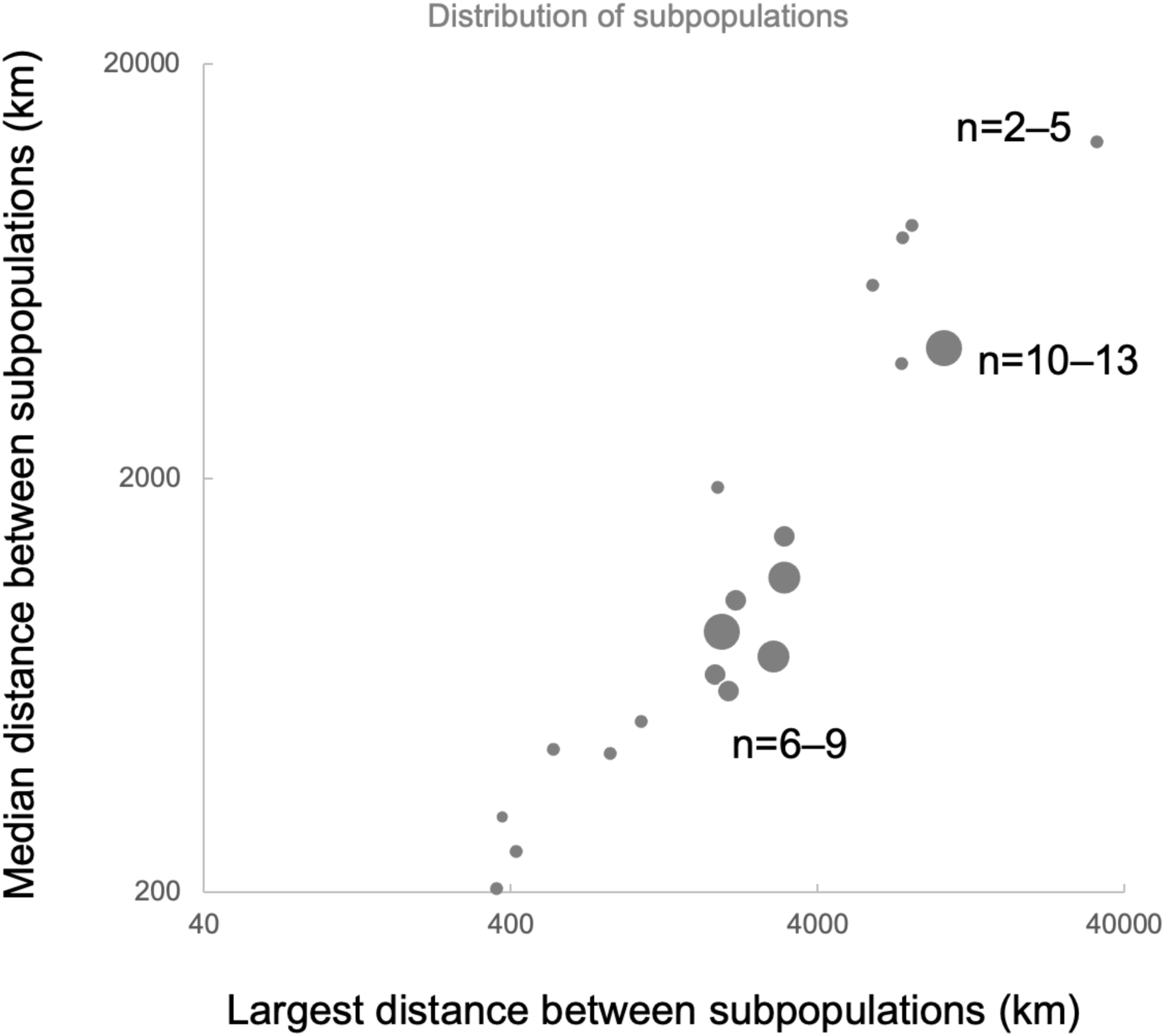
Covariation among three of the sampling variables in all 20 studies of phylogeography in sea slugs. Three symbol sizes show differences in the number of subpopulations (n). Note logarithmic scale for distance.

Pairwise differentiation between subpopulations was broadly distributed from *Φ* _ST_ < 0 (no differentiation) to *Φ* _ST_ ∼1 (fixed differences; Figure 2). The maximum possible value of *Φ* _ST_ in any system may be << 1 and depends on the variability of the genetic marker and the extent of variation within subpopulations (e.g., Alcala and Rosenberg 2019), so interpreting quantitative variation in *Φ* _ST_ values among studies can be challenging, but relatively high *Φ* _ST_ values are commonly interpreted as evidence for some combination of restricted gene flow, small effective population size, and recent metapopulation dynamics, whereas lower *Φ* _ST_ values are commonly interpreted as evidence of substantial gene flow (such as gene flow by dispersing planktonic veligers of marine gastropods). Most subpopulation pairs were <1000 km apart (Figure 2), with *Φ* _ST_ values less than 0.2 (and many of those values were not significantly different from zero in the published AMOVA). Even among subpopulation pairs >4000 km apart, *Φ* _ST_ varied from ∼0 to ∼0.7 (and statistically significant). Median *Φ* _ST_ varied from <0.005 (in a pulmonate) to >0.5 (in several nudibranchs, dorids, and sacoglossans). Similarly, the smallest distance between subpopulations with *Φ* _ST_ > 0 varied from ∼80 to ∼8000 km.

**Figure 2.**
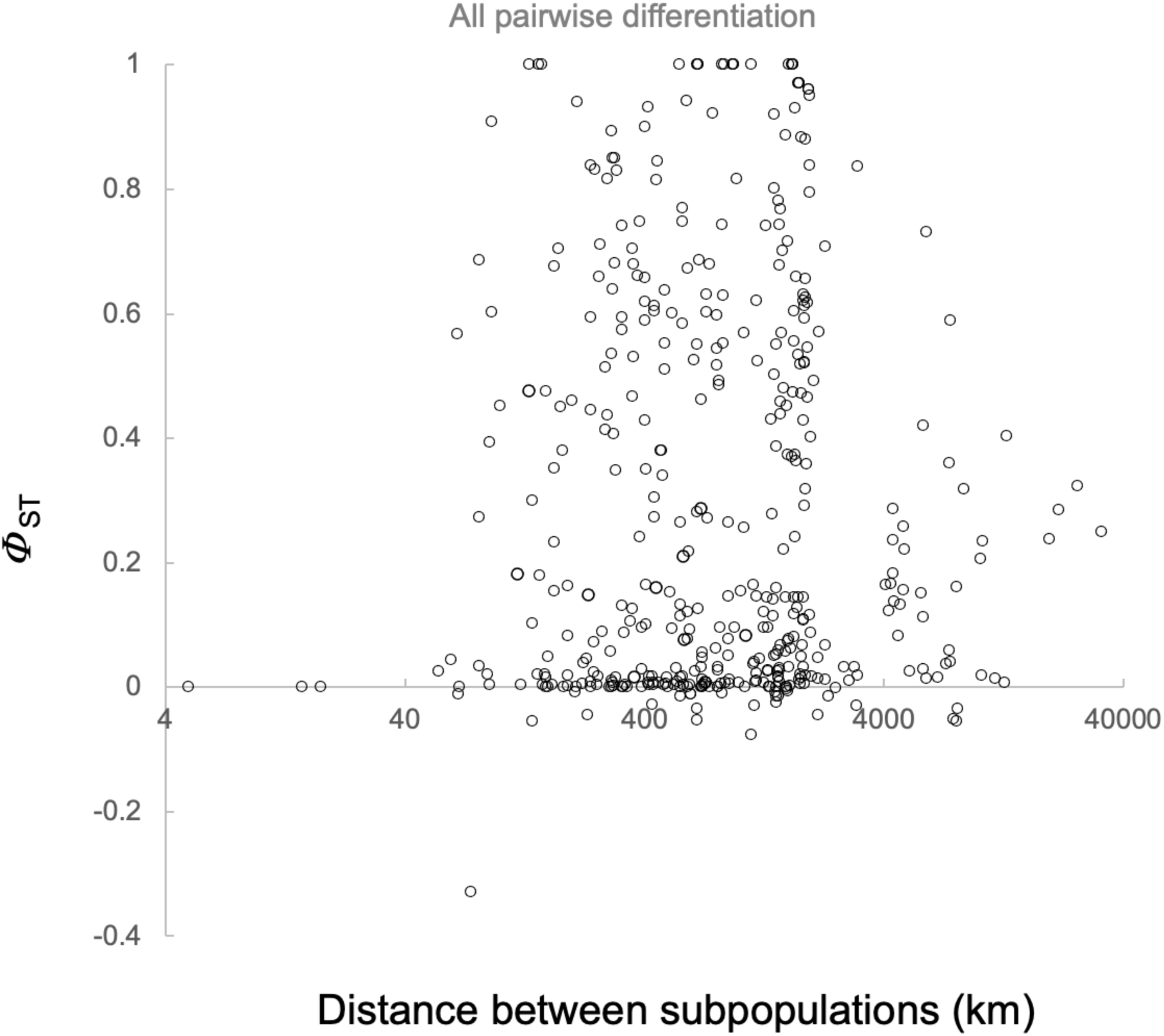
Variation in pairwise differentiation and pairwise distance among all subpopulations in all 20 studies of phylogeography in sea slugs. Note logarithmic scale for distance.

In a principal components analysis of the six variables shown in Table 1, the two measures of the size of the sample (number of subpopulations, number of individuals) were positively correlated across studies (*r* = 0.43; Table 2), as were the two measures of the geographical distribution of samples (median distance among all subpopulations, largest distance among subpopulations (*r* = 0.89; see Figure 1). Those correlations may reflect in part the costs or logistical tradeoffs between large-scale breadth and local intensity or granularity of sampling subpopulations, and researcher expectations about the geographical scale of genetic variation. By contrast, the two population genetic variables (median *Φ* _ST_, smallest distance between subpopulations with *Φ* _ST_ > 0) were mostly independent of each other (*r* = -0.07; Table 2). However, each was associated with a combination of several sampling variables.

**Table 2.**
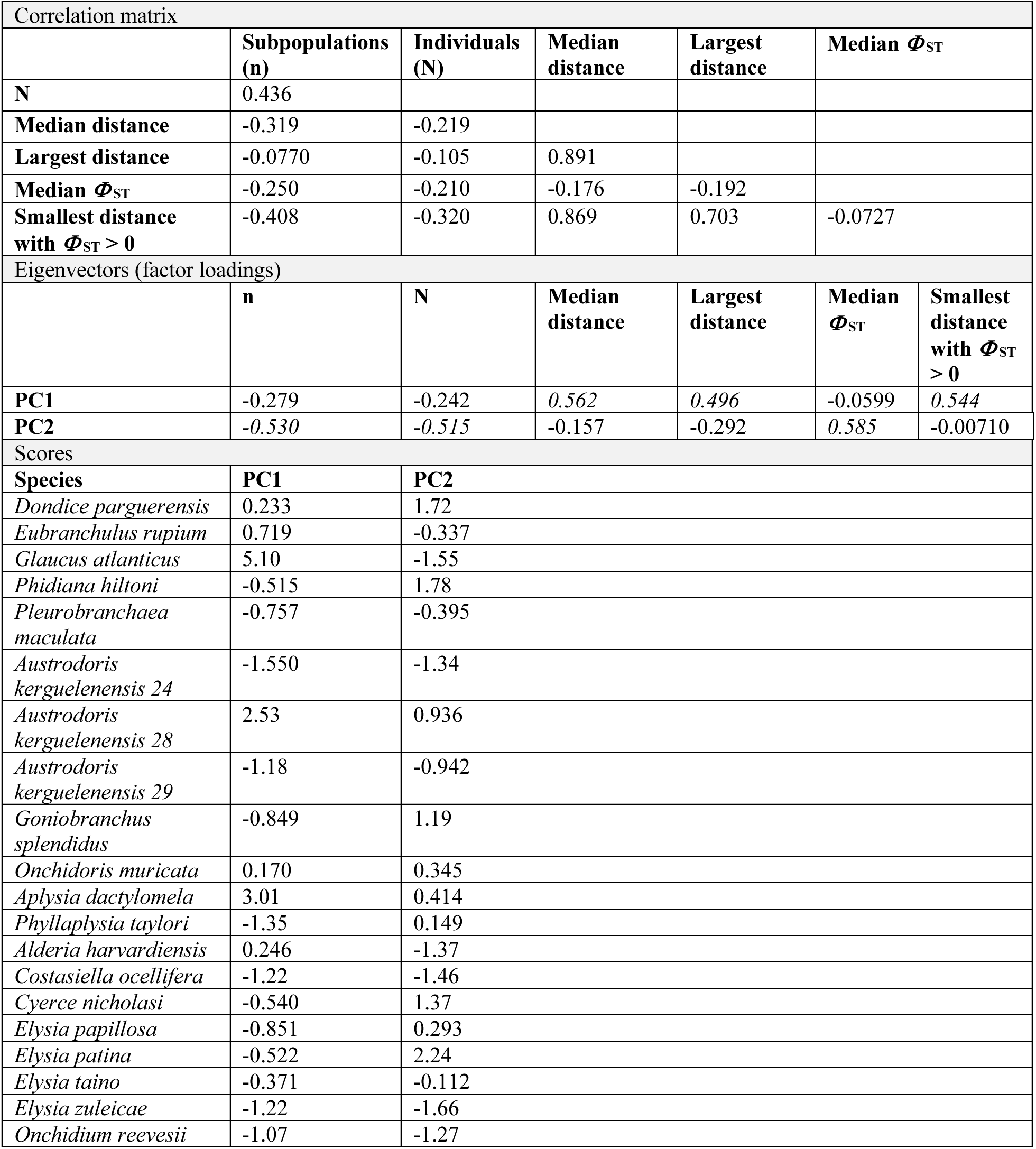
Results from a principal components analysis of the six sampling and population genetic variables in 20 studies of sea slugs in Table 1. Only scores for the first and second principal components are shown here; see Appendix 3 for full results (script, standardized variables, all eigenvalues and eigenvectors). *Factor loadings with contrasting large absolute values are shown in italics*.

| Correlation matrix |  |  |  |  |  |  |
|---|---|---|---|---|---|---|
|  | **Subpopulations (n)** | **Individuals (N)** | **Median distance** | **Largest distance** | **Median *Φ*<sub>ST</sub>** |  |
| **N** | 0.436 |  |  |  |  |  |
| **Median distance** | -0.319 | -0.219 |  |  |  |  |
| **Largest distance** | -0.0770 | -0.105 | 0.891 |  |  |  |
| **Median *Φ*<sub>ST</sub>** | -0.250 | -0.210 | -0.176 | -0.192 |  |  |
| **Smallest distance with *Φ*<sub>ST</sub> > 0** | -0.408 | -0.320 | 0.869 | 0.703 | -0.0727 |  |
| Eigenvectors (factor loadings) |  |  |  |  |  |  |
|  | **n** | **N** | **Median distance** | **Largest distance** | **Median *Φ*<sub>ST</sub>** | **Smallest distance with *Φ*<sub>ST</sub> > 0** |
| **PC1** | -0.279 | -0.242 | *0.562* | *0.496* | -0.0599 | *0.544* |
| **PC2** | *-0.530* | *-0.515* | -0.157 | -0.292 | *0.585* | -0.00710 |
| Scores |  |  |  |  |  |  |
| **Species** | **PC1** | **PC2** |  |  |  |  |
| *Dondice parguerensis* | 0.233 | 1.72 |  |  |  |  |
| *Eubranchulus rupium* | 0.719 | -0.337 |  |  |  |  |
| *Glaucus atlanticus* | 5.10 | -1.55 |  |  |  |  |
| *Phidiana hiltoni* | -0.515 | 1.78 |  |  |  |  |
| *Pleurobranchaea maculata* | -0.757 | -0.395 |  |  |  |  |
| *Austrodoris kerguelenensis 24* | -1.550 | -1.34 |  |  |  |  |
| *Austrodoris kerguelenensis 28* | 2.53 | 0.936 |  |  |  |  |
| *Austrodoris kerguelenensis 29* | -1.18 | -0.942 |  |  |  |  |
| *Goniobranchus splendidus* | -0.849 | 1.19 |  |  |  |  |
| *Onchidoris muricata* | 0.170 | 0.345 |  |  |  |  |
| *Aplysia dactylomela* | 3.01 | 0.414 |  |  |  |  |
| *Phyllaplysia taylori* | -1.35 | 0.149 |  |  |  |  |
| *Alderia harvardiensis* | 0.246 | -1.37 |  |  |  |  |
| *Costasiella ocellifera* | -1.22 | -1.46 |  |  |  |  |
| *Cyerce nicholasi* | -0.540 | 1.37 |  |  |  |  |
| *Elysia papillosa* | -0.851 | 0.293 |  |  |  |  |
| *Elysia patina* | -0.522 | 2.24 |  |  |  |  |
| *Elysia taino* | -0.371 | -0.112 |  |  |  |  |
| *Elysia zuleicae* | -1.22 | -1.66 |  |  |  |  |
| *Onchidium reevesii* | -1.07 | -1.27 |  |  |  |  |

The first and second principal components accounted for 48% and 26% of the variance in standardized values. Scores on PC1 (Table 2) represented mainly a contrast between the geographical distribution of samples (median distance and largest distance between subpopulations, with coefficients 0.56 and 0.49) and one of the population genetic variables (smallest distance with a significant *Φ* _ST_ value, or the granularity of genetic differentiation, coefficient 0.54). Studies with high scores on PC1 had coarse-scale genetic differentiation (on large spatial scales only) among samples separated by large distances. Scores on PC2 represented mainly a contrast between sample size (the number of subpopulations and individuals, with coefficients -0.53 and -0.51) and the other population genetic variable (median *Φ* _ST_ value, coefficient 0.58). Studies with high scores on PC2 had strong differentiation among most subpopulations with relatively small samples of subpopulations or individuals.

In that two-dimensional space, a few studies and species stood out (Figure 3). In their analysis of the pelagic nudibranch *Glaucus atlanticus*, Churchill et al. (2014) sampled few regional subpopulations on the scale of oceanic basins (∼10^4^ km) and found modest *Φ* _ST_ values (∼0.2) on that coarse spatial scale with a few significantly differentiated subpopulation pairs (≥ 8700 km apart; Table 1). The two other studies with high PC1 scores in the dorid *Austrodoris kerguelenensis* species 28 (Maroni and Wilson 2022) and the sea hare *Aplysia dactylomela* (Valdés et al. 2013) each sampled only two regional subpopulations separated by > 7500 km (Table 1).

**Figure 3.**
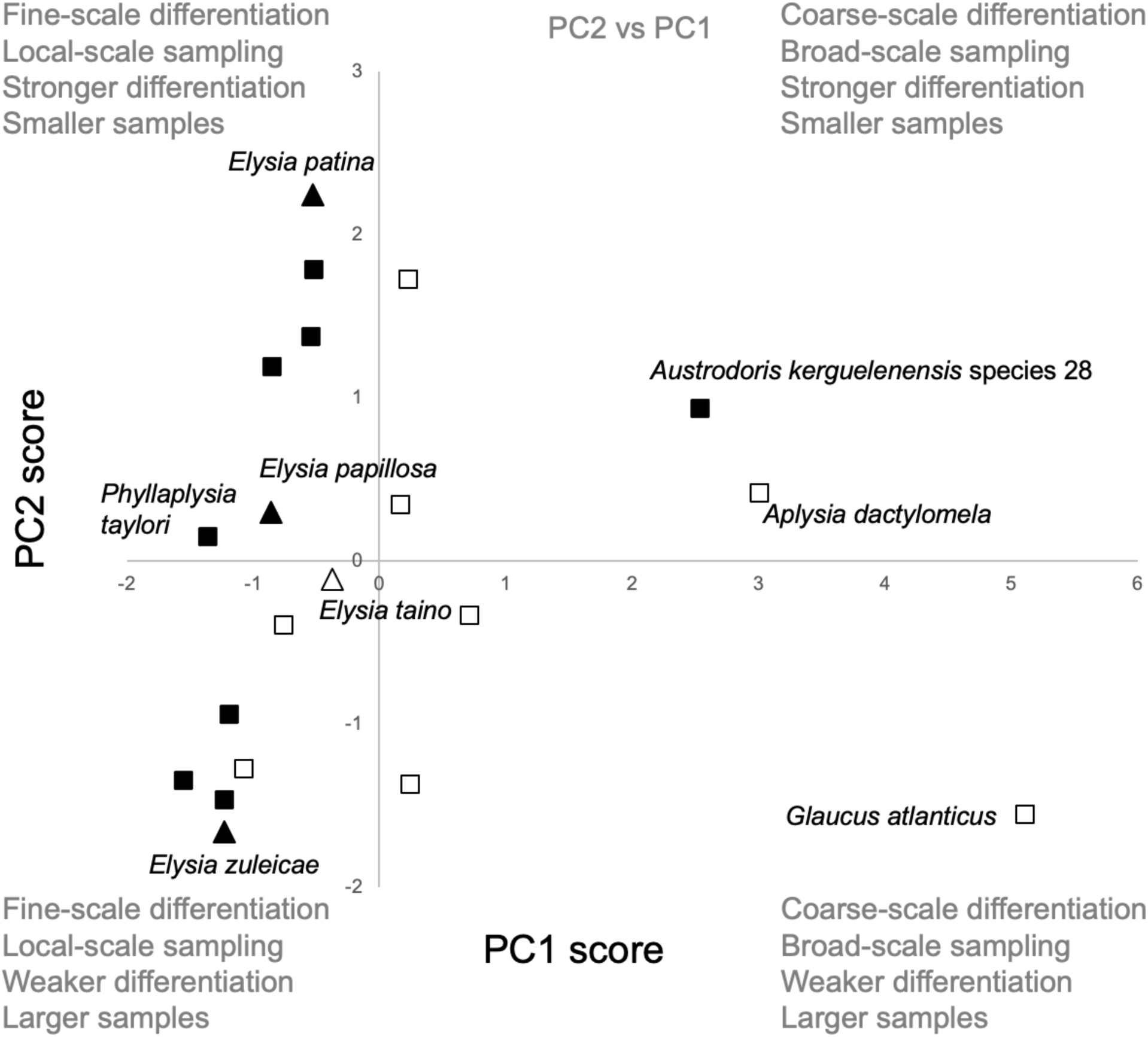
Scores on the 1^st^ and 2^nd^ principal components from an analysis of six sampling and population genetic variables in 20 studies of phylogeography in sea slugs (data in Table 1; PCA results in Table 2; see Appendix 3). PC1 represents a spatial contrast: studies with high scores sampled subpopulations on large spatial scales and found significant *Φ* _ST_ values only between subpopulations separated by large distances. PC2 represents a contrast between sample size and magnitude of differentiation: studies with high scores had smaller samples and larger *Φ* _ST_ values. Scores for nine species with exclusively feeding planktonic veliger larvae are shown as open symbols. Scores for four species of *Elysia* are shown as triangles.

Most other studies had similar low PC1 scores (studies carried out over smaller geographical distances, with statistically significant differentiation on finer spatial scales <2000 km) and differed along the second PC axis. At the two extremes, studies of several closely related species of sacoglossans by Krug et al. (2026) discovered high median *Φ* _ST_ values among a few samples from Florida and the Bahamas (*Elysia patina*; Figure 3) or much lower median *Φ* _ST_ values from many samples across the Caribbean and western Atlantic Ocean (*E. zuleicae*).

The choice of a sampling design (many or few samples separated by large or small distances) to detect strong or weak differentiation on coarse or fine scales might depend on the mode of larval development. Among the nine studies of species with exclusively planktotrophic veliger larvae (Table 1), median score on PC1 was higher (0.23) and median score on PC2 was lower (-0.33) than median scores among eleven other studies (-0.85, 0.29) (Figure 3). Although there was considerable variation within each of those groups of studies (e.g., *Austrodoris kerguelenensis* species 28; Maroni and Wilson 2022), those differences between medians suggest that studies of planktotrophic species tended to collect larger samples of subpopulations separated by larger distances, and tended to discover lower *Φ* _ST_ values that were significantly different from zero only on more coarse geographical scales.

Other contrasts between taxonomically close groups in our review were consistent with those differences. For example, in their broad and intensive sampling of the sacoglossan *Costasiella ocellifera* with subpopulation differences in the mode of reproduction, Ellingson and Krug (2015) found a bimodal distribution of pairwise *Φ* _ST_ values (Figure 4), including a larger proportion of values >0.4 for comparisons in which one or both subpopulations had direct development of benthic embryos, and a larger proportion of values <0.2 for comparisons in which both subpopulations had planktonic veliger larvae. The two studies of sea hares (Louie 2003; Valdés et al. 2013) with different modes of larval development also had contrasting sampling designs and patterns of genetic differentiation (Table 1; Figure 3).

**Figure 4.**
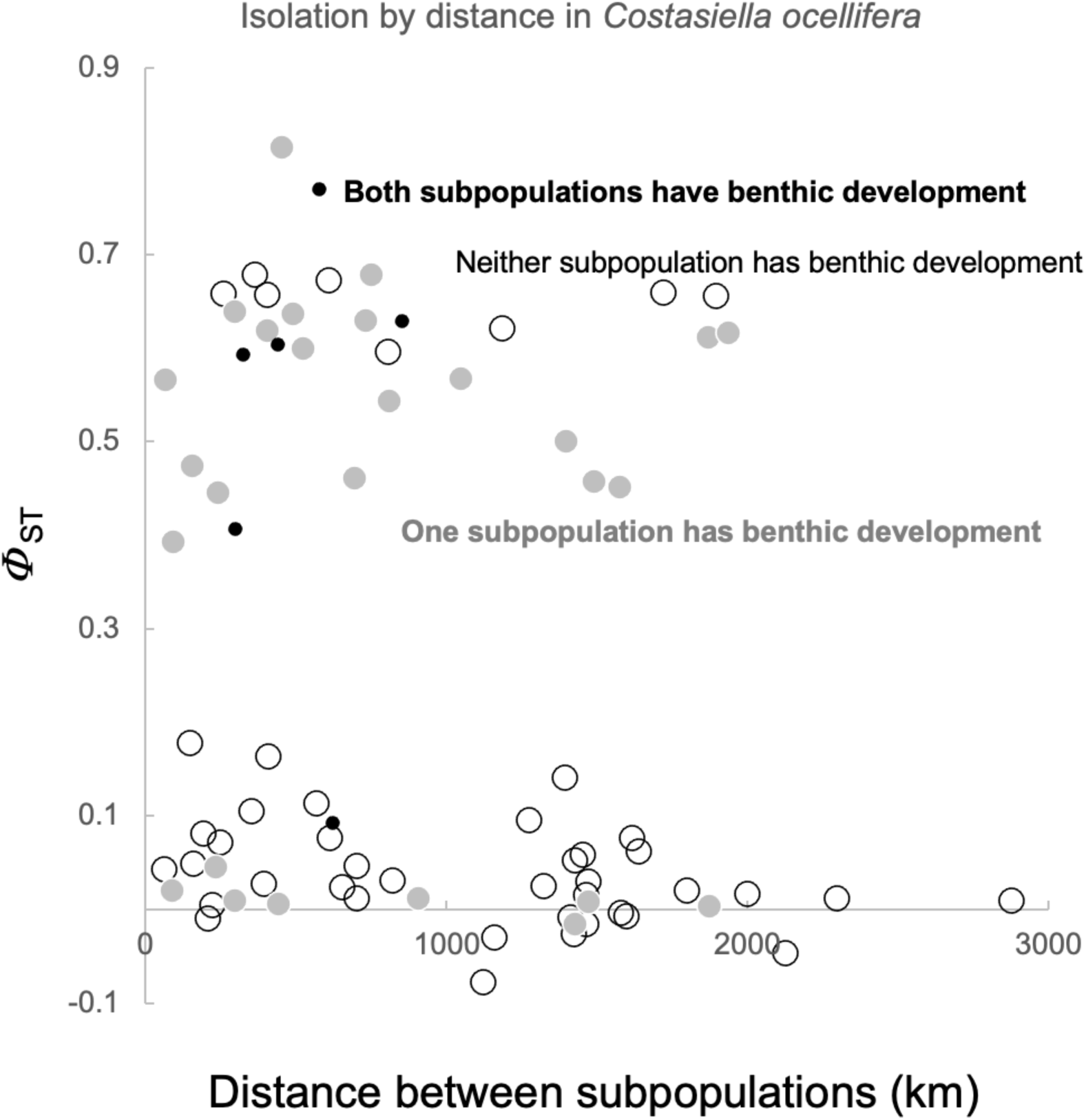
Bimodal distribution of *Φ* _ST_ values among pairs of subpopulations of *Costasiella ocellifera* (from Ellingson and Krug 2015) that have similar or different modes of larval development.

## DISCUSSION

We reviewed studies of “phylogeography unplugged” in nudibranchs and other sea slugs and found wide variation in sampling and in the magnitude or geographical scale of genetic differentiation. Correlations among sampling variables probably reflect a combination of natural features of the sea slugs (their geographical range and abundance) and tradeoffs or constraints on sampling design for studying their population genetic differentiation. In the clustering analysis, we found coarse-grained spatial structure in studies with samples collected on large spatial scales (PC1), and stronger differentiation in studies with small samples of individuals and subpopulations (PC2). Studies of planktotrophic species used larger samples collected on larger spatial scales, and found weaker differentiation among distant subpopulations (Todd et al. 1998; Grosberg and Cunningham 2001; Krug et al. 2015). Those results are broadly similar to patterns of spatial genetic variation and connectivity associated with sample sizes, the geographical scale of sampling, and the mode of larval development in marine fishes and in other marine invertebrates documented using regression methods (Riginos et al. 2011; Selkoe and Toonen 2011) rather than ordination or clustering methods (PCA; Figure 3).

The associations represented by PC1 seem obvious: Studies that sampled only a few widely separated subpopulations could find only the most coarse-grained differentiation (on the order of ∼10,000 km). Other studies were carried out over smaller distances and found significant differentiation on less coarse spatial scales (<2000 km apart). Some researchers in the studies we reviewed may have correctly predicted the occurrence of differentiation only on coarse spatial scales (e.g., Churchill et al. 2014), assumed that subpopulations are not differentiated on smaller scales, and adapted their sampling design accordingly. Sampling in such studies (e.g., *Aplysia dactylomela*; Valdés et al. 2013) may deliberately include fewer subpopulations all separated by large distances (high scores on PC1). Other studies of organisms with similar high larval dispersal potential carried out on a smaller spatial scale (lower score on PC1, e.g., *Elysia taino*; Krug et al. 2026) but with similar median *Φ* _ST_ value (similar score on PC2) may have erred on the side of sampling a larger number of subpopulations (separated from each other by smaller distances) than was needed to capture the expected spatial variation, genetic diversity, and coalescent history. Such studies may have benefitted in other ways from that more intensive sampling on smaller spatial scales. For example, Krug et al. “predicted little structure in *E. taino*, in which all sampled demes produced planktotrophic offspring expected to drive high connectivity”, but the authors sampled many localities that included *E. taino* on a relatively small spatial scale in several parts of the Caribbean Sea because of previous concerns about the taxonomic validity of *E. taino* and because it is broadly sympatric with and morphologically similar to its sister species *E. papillosa* (Krug et al. 2026). Most other authors of the studies we reviewed did not explicitly explain or justify their sampling designs in these ways, but the associations we observed (Figure 1) suggest that tradeoffs or resource limitations combined with expectations about the geographical scale of differentiation together influence observed variation in sampling design, and may account for the association between sampling variables and the population genetic variables.

The associations represented by PC2 may have a similar interpretation. Among the five studies with the highest scores on PC2 (Figure 3), with large median *Φ* _ST_ values from small samples (García-Méndez et al. 2022; King et al. 2019; Winters et al. 2017; Moreno 2024; Krug et al. 2026), four were studies of species with a short planktonic veliger stage in the life cycle and limited potential for larval dispersal and gene flow (*Goniobranchus splendidus, Cyerce nicholasi, Phidiana hiltoni, Elysia patina*; Figure 3). In those four studies, the geographical scale was smaller (<1000 km) and the smallest distance between subpopulations with statistically significant differentiation was much smaller (100–300 km; Table 1) than most other studies.

Similarly, among other studies with the lowest scores on PC2, each with a low median *Φ* _ST_ value in a larger sample, many were also studies of species in which individuals in some or all subpopulations have modes of reproduction with limited potential for dispersal and gene flow (e.g., *E. zuleicae*; Figure 3; Krug et al. 2026). One interpretation of that pattern is that for some studies (with low scores on PC1 on smaller spatial scales), a relatively small sample could successfully capture existing genetic diversity and differentiation (high scores on PC2), and that more intensive (over)sampling in other studies relative to the dispersal potential of the organisms may have led to the discovery of more haplotype sharing among pairs of subpopulations and a larger number of relatively lower pairwise *Φ* _ST_ values (low scores on PC2).

Can other researchers use those patterns to design successful sampling in the absence of an unplugged survey across the species range? If accurate expectations about population differentiation caused the covariation between sampling and *Φ* _ST_ among studies (e.g., some studies with high PC2 scores focused on species expected to show strong differentiation, and the authors correctly designed those studies with less intensive sampling of subpopulations and individuals; Figure 3), then future studies could similarly adopt sampling designs that will capture a large proportion of expected spatial variation given some prior knowledge based on the biology of the organisms. In particular, future genome-wide studies of species with planktotrophic veligers and expected high potential for dispersal and gene flow might emphasize smaller sample sizes (PC2) and larger geographical scales (PC1) like those in Figure 1 to capture genetic variation that is expected to be of smaller magnitude overall and detectable only on relatively coarse scales. Studies of species with less potential for planktonic larval dispersal might adequately capture genetic variation with larger samples collected on smaller geographical scales. However, the specific meaning of “large samples” or “small scales” might be subjective and differ among study systems, and might be influenced by many other considerations within a single study.

## ACKNOWLEDGMENTS

We thank several colleagues who kindly provided links to articles or datasets (I. Ekimova, D. Grishina, D. Jacobs, P. Krug, R. Väinölä, Á. Valdés) or gave us helpful suggestions on an early version of the manuscript (P. Marko). We are supported by grants and scholarships from the Natural Sciences and Engineering Research Council of Canada and by Simon Fraser University.

## List of appendices

**Appendix 1** List of articles from the literature review with inclusion/exclusion criteria and notes.

**Appendix 2** Pairwise distances and *Φ* _ST_ values among all subpopulation pairs in all 20 studies.

**Appendix 3** Principal components analysis (results, code).

